# Identification of the bona fide active center of influenza A virus polymerase acidic protein as the antiviral target

**DOI:** 10.1101/2025.08.06.669014

**Authors:** Shan Xu, Xiaoyang Wang, Lihong Zhang, Yanli Tu, Yueqi Wu, Siying Du, Wei Wang, Shan Xu, Yigang Tong

**Author notes:** Corresponding author. (Y.G.T.); (S.X.); (W.W.). These authors contributed equally to this work (Shan Xu, Xiaoyang Wang, Lihong Zhang, Yanli Tu).

## Abstract

The influenza virus PA protein, a core subunit of RNA polymerase, is critical for viral replication and a key antiviral target. Contrary to the prevailing view that its endonuclease active site resides in the N-terminal domain (PAn, residues 1-319), we identified a C-terminal truncation (PAc, 320-716) of H1N1 PA that retained high endonuclease activity, while PAn showed minimal activity. Crystal structural analysis revealed that PAc binds substrate RNA via the YDS motif (residues 393-395), with mutations abolishing RNA association. Surprisingly, baloxavir acid (BXA), a known PA inhibitor, specifically targeted PAc rather than PAn, supported by molecular docking showing higher binding affinity to PAc. Mutations at PAc-BXA interaction sites identified F707A as a potential drug resistance mutation. Virtual screening targeting PAc identified methotrexate as potent inhibitors (EC_50_ < 100 nM), which effectively suppressed viral replication in vitro and alleviated symptoms in influenza-infected mice. Our findings redefine PA’s catalytic architecture and establish PAc as a novel platform for antiviral discovery, offering strategies to combat drug resistance through structure-guided inhibitor design.

## Introduction

Influenza, a respiratory infectious disease caused by the influenza virus, is among the greatest global public health challenges. According to statistics, approximately 1 billion people worldwide are infected with the influenza virus annually, resulting in 290,000-650,000 deaths (*1*). Influenza viruses belong to the *Orthomyxoviridae* family, and comprise four types: A, B, C, and D. Influenza A virus is highly infectious, and its antigens can mutate easily, leading to several world pandemics (*2*). Vaccination and drug treatment are the main strategies to prevent and treat influenza infection. Based on the various epitopes of the influenza virus, vaccines can effectively prevent infection with multiple influenza subtypes (*3*).However, owing to the long production cycle, strict storage conditions and high cost, the supply of influenza vaccines is generally insufficient to meet the need for timely emergency response (*4*).Especially, People tend to prefer drug treatment rather than vaccination against influenza, which increases the importance of drug development (*5*).

Anti-influenza drugs inhibit influenza virus replication by targeting the key steps in the virus life cycle (*6*). Influenza virus enters the host cell via hemagglutinin (HA)-mediated endocytosis. The acidic environment of the lysosome induces membrane fusion with the participation of the M protein, resulting in the release of the virus genome into the cytoplasm. Viral RNA (vRNA) and messenger RNA (mRNA) are produced by the RNA-dependent RNA polymerase (RdRp) complex (PB1, PB2, and PA proteins) with RNA templates. mRNA is translated into viral proteins, which assemble with vRNA into progeny virions. To successfully release the new virus, neuraminidase (NA) cleaves sialic acids and disrupts the interaction between HA and sialic acid receptors. Currently, four first-line anti-influenza drugs approved by the U.S. Food and Drug Administration are recommended by the Centers for Disease Control and Prevention. Peramivir, zanamivir, and oseltamivir are NA inhibitors, which suppress influenza infection by hindering virus release (*7*). Baloxavir marboxil (BXM) depresses the endonuclease activity of PA, destroying the functioning of the RdRp complex (*8*). Despite the high antiviral efficiency of these medicines, rapid mutation of the influenza virus genome contributes to the risk of drug resistance (*9, 10*). Therefore, novel anti-influenza drugs should be designed to combat new potential variants.

Novel drug targets provide the basis for the development of new antiviral drugs. For instance, S31N mutation in M2 generates new RNA release channels, which leads to amantadine resistance (*11*). Such mutation-dependent channels could be a novel target in the design of drugs against new influenza variants. Traditionally, each component of RdRp was used as a target for designing compounds to inhibit the activity of a certain protein, thereby inhibiting the functioning of the RdRp complex (*12, 13*). Recently, the interaction interface of RdRp has been identified as a potential antiviral drug target. For example, compounds have been designed to inhibit the interaction between the C-terminus of the PA protein and the N-terminus of the PB1 protein, disrupting RdRp assembly and thereby inhibiting viral replication (*14, 15*). In the present work, we have accidentally discovered a novel PA truncation (PAc) of influenza A virus, which showed remarkably high endonuclease activity. Furthermore, the PAc domain was demonstrated to be a bona fide target for antiviral drugs, such as BXA. The identification of PAc as the active center of PA suggests a new mechanism of viral RNA amplification. More importantly, this provides novel insights for anti-influenza compound screening.

## Results

### Identification of a PA endonuclease C-terminal truncation (PAc)

To explore the enzyme activity of PA, which is vital for influenza virus RNA replication, we expressed the influenza A PA protein in *Escherichiacoli*. Surprisingly, we found that a fragment stably existed in the cell lysate when PA was expressed at 37 °C (Fig. 1A). By contrast, the specific fragment was not obvious at 16 °C. Western blot was used to detect the induced bacterial lysate targeting the C-terminal of the PA protein. The results indicated that the full-length PA was successfully induced at both temperatures. The specific smaller band at 37 °C seemed to be a PA fragment at the C-terminal. To clarify the amino acid sequence, we incised the band and performed N-/C-terminal sequencing using liquid chromatography–mass spectrometry (Fig. 1B).

**Fig. 1.**
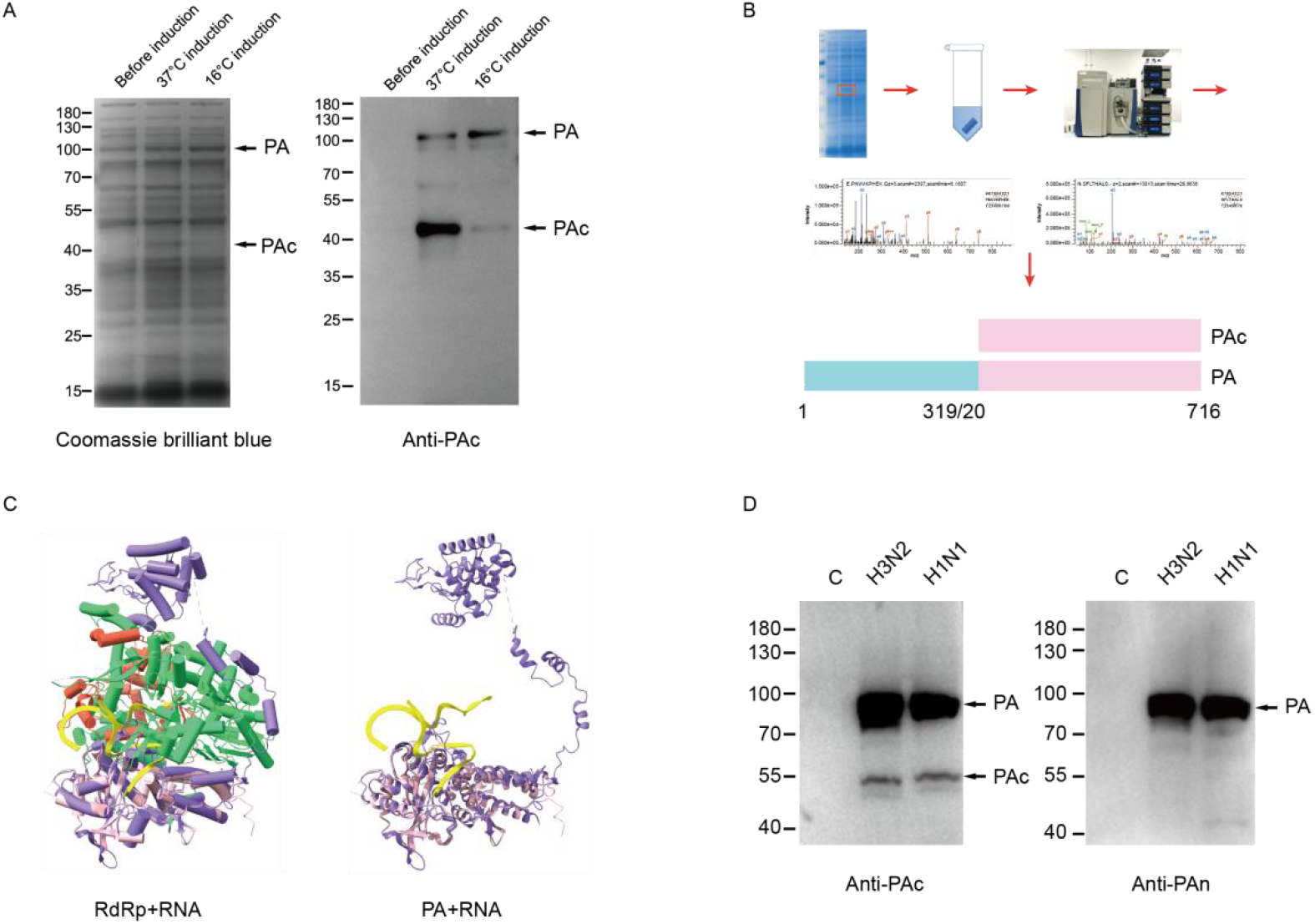
Identification of novel influenza PA C-terminal truncation. A. Induced expression of influenza H1N1 PA protein at 37°C or 16°C in *E*.*coli*. Coomassie brilliant blue staining and western blot were used to detect the protein in cell lysates. B. The N/C terminal sequence of PAc was determined by LC-MS/MS. The specific band of PAc from SDS-PAGE was cut out for protein sequencing. Sequencing result was indicated in the schematic diagram. C. The structure of H17N10 RdRp complex with RNA (PDB:6t0n), in which PB1 in green color, PB2 in red color, PA in purple color and RNA in yellow color. H1N1 PA C-terminal domain structure (PDB:2znl) was aligned to the complex in pink color. D. MDCK cell lysate was detected with western blot by using different antibodies that target influenza PA N-terminal or PA C-terminal domain. MDCK cell were infected with influenza virus H3N2 or H1N1 for 24h before the detection of the cell lysate.

The fragment was demonstrated to be the C-terminal truncation of PA (PAc) between amino acids 320 and 716. The identification of PAc prompted us to investigate its function and physiological importance. Structure determination of the H17N10 RdRp complex with RNA provided some hints (*16,17*) (Fig.1C). The H1N1 PA structure was aligned to the H17N10 RdRp structure and showed a similar structure to that of H17N10 PA (*18*). Importantly, the RNA substrate was closely associated to PAc rather than PAn, indicating the role of PAc as the active center. In addition, H1N1- and H3N2-infected cell lysate were detected using western blot with an antibody targeting the N-terminal or C-terminal PA (Fig. 1D). As shown in the figure, PAc was present in cells infected with influenza A virus. Moreover, immunofluorescence staining targeting the N-terminal or C-terminal domain of PA in influenza-infected cells was performed. The results revealed the specific subcellular localization and potential physiological function of PAc (Fig. S1).

### PAc is the catalytic domain of PA with high endonuclease activity

To test the endonuclease activity of the identified PAc, full-length PA, PAc, and PAn expression vectors were designed and the expressed proteins were purified. The purified proteins stained using Coomassie blue were consistent with the theoretical size as further proved by western blot analysis (Fig. 2A and B). An in vitro nucleic acid degradation assay was used to detect the endonuclease activity of full-length and truncated PA (*19, 20*), and the quantitative results were based on three independent repetitions (Fig. 2C). As shown, PAc and PA exhibited high endonuclease activity in degrading RNA, whereas the enzyme activity of PAn was significantly lower. A FRET (Fluorescence Resonance Energy Transfer) -based assay using FAM/BHQ1-labeled RNA also demonstrated that the endonuclease activity of PAc was much higher than PAn (*21*) (Fig. 2D). PAn 1-209 that reported to show endonuclease activity cleaved RNA substrates only at elevated protein concentrations with extended reaction times (Fig. S2). These results indicated that PAc might be the bona fide catalytic domain of PA. To confirm this, the structure of the PA–RNA complex was analyzed to identify amino acids that are likely to be vital for the interaction between PA and the RNA substrate (Fig. 2E). We obtained various mutants and detected their endonuclease activity (Fig. 2F,and Fig. S3). It was demonstrated that amino acids 393–395 in H1N1 PA are indispensable for PA endonuclease activity. And the quantitative data were based on three independent repetitions (Fig. 2G). The vital role of 393–395 residues, which reside in the PAc domain, further proved PAc to be the catalytic domain of PA.

**Fig. 2.**
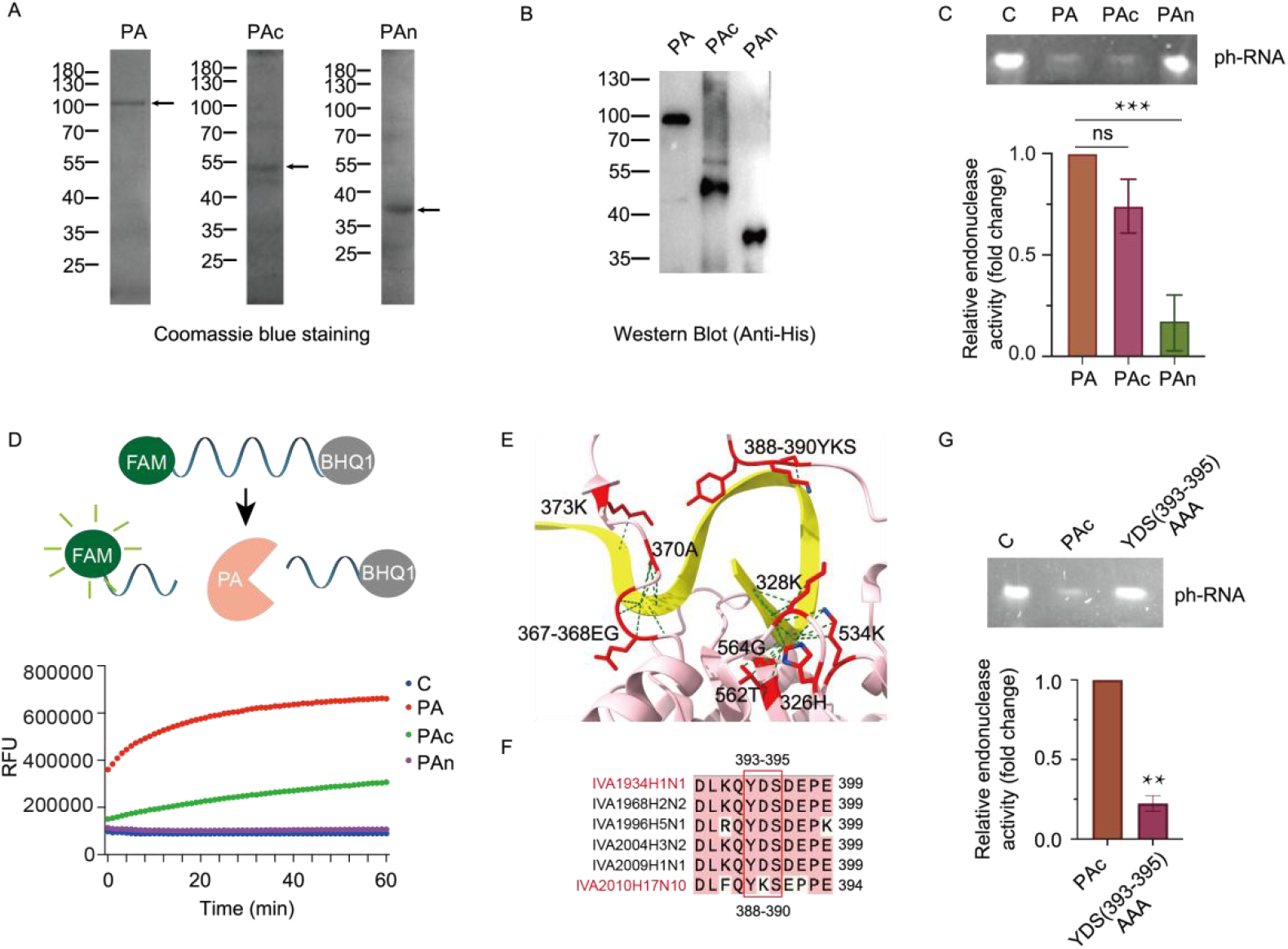
Newly identified influenza PAc domain show high endonuclease activity. A. Expression and purification of influenza PA, PAc and PAn in *E*.*coli*. The purified proteins were subjected to SDS-PAGE and visualized by Coomassie Brilliant Blue staining. The targeted bands are indicated with arrows. B. The purified PA, PAn and PAc were detected by western blot using anti-His tag antibody. C. Upper panel: Endonuclease activity of PA, PAc and PAn with RNA as the substrate. 750nM ph-RNA substrate was co-incubated with 300nM proteins in reaction buffer solution for 10 minutes at 37°C. Reaction products were analyzed by 8% urea-PAGE and visualized with YeaRed nucleic acid stain. Lower panel: Relative endonuclease activity of PA, PAc and PAn was quantified based on three replications in upper panel. ***: p < 0.001. Reaction product bands were quantitatively analyzed with Image J software to determine the relative enzymatic activities of different proteins. D. Upper panel: A schematic representation of the endonuclease activity of the PA protein was analyzed using the FRET principle. A FAM-BHQ1 quenched RNA probe exhibited no detectable fluorescence. Upon PA protein cleavage, FAM separation from BHQ1 restored fluorescence, enabling measurement of PA’s endonuclease activity via fluorescence intensity. Lower panel: The FRET-based assay was used to detect the endonuclease activities of PA, PAc and PAn with FAM/BHQ1-labeled RNA as the substrate. E. Detailed structure of H17N10 PA with RNA. Predicted residues responsible for the interaction between PA and RNA are highlighted in red. RNA was in yellow color and PA protein was in pink color. F. Amino acid alignment of H17N10 and several human influenza A PA. YKS (388-390) amino acids in H17N10 PA correspond to YDS (393-395) in H1N1 PA. G. Upper panel: Endonuclease activity of H1N1 wild-type PAc and YDS(393-395)AAA mutant. 750nM ph-RNA substrate was co-incubated with equal amounts of wild-type PAc and mutant for 10 minutes at 37°C. Lower panel: Relative endonuclease activity H1N1 wild-type PAc and YDS(393-395)AAA mutant was quantified based on the results of three replicates in upper panel. Image J software was used to analyze the product bands to quantify the enzyme activity. **: p < 0.01 vs wild-type PAc.

### PAc domain is the target of a first-in-class anti-influenza drug (baloxavir acid)

Identification of the catalytic function of PAc indicates its potential as an antiviral drug target. Baloxavir acid (BXA) is derived from the prodrug BXM, which inhibits the replication of influenza virus by depressing PA endonuclease activity (21). To investigate whether BXA and BXM target the PAc domain, various concentrations of BXA and BXM were applied in the RNA degradation system (Fig. 3A and B, and Fig. S4). The quantified results were based on three independent repetitions. The enzyme activity of PA was inhibited by BXA in a dose-dependent manner (Fig. 3A). The inhibition of PAc activity by BXA was also concentration-dependent (Fig. 3B). The results demonstrated that BXA directly targets PAc and can inhibit endonuclease activity. To further validate this finding, we explored the BXA binding sites in PAc. As indicated from the results of molecular docking, BXA binds to residues 616, 634, 638, 661, 701, 702, and 705 of H17N10 PA (Fig. 3C). When aligned with H1N1, these amino acid sites correspond to 621, 639, 643, 666, 706, 707, and 710 (Fig. 3D). Except for 707, mutations of all other residues suppressed the endonuclease activity of H1N1 PA itself, which made it difficult to explore its interaction with BXA (Fig. 3E and Fig. S5). Instead, the 707 mutant showed similar enzyme activity to wild-type PA but clearly impaired the inhibitory effect of BXA on PA activity (Fig. 3F and G). These results demonstrated that BXA directly targets the PAc domain, and the 707 site is a predominant factor in BXA binding and function.

**Fig. 3.**
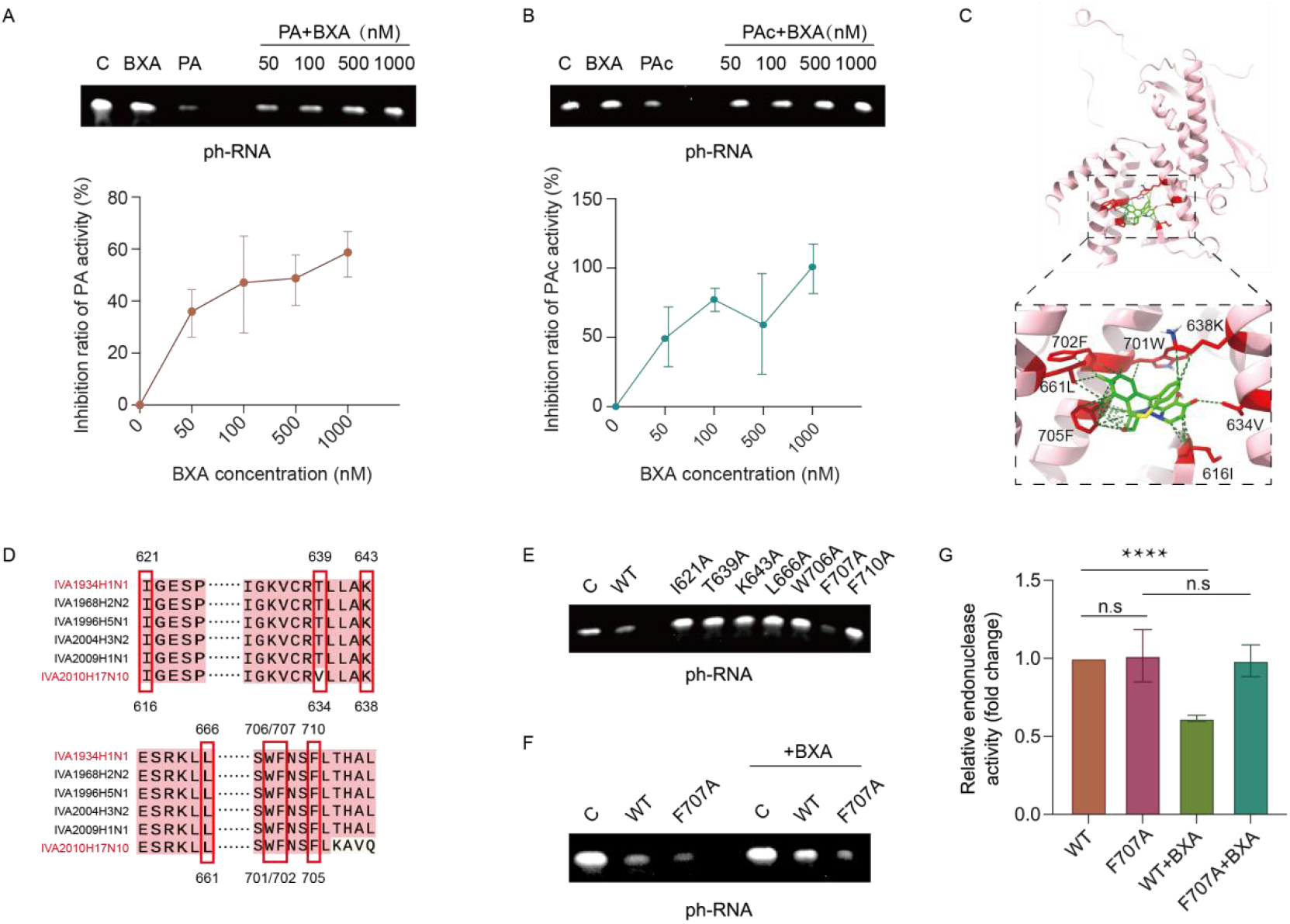
Influenza PAc domain is the drug target of Baloxavir acid (BXA). A. The inhibitory effect of various concentrations of BXA on endonuclease activity of H1N1 PA. The results were quantified from three replicates. Various concentrations of BXA and 300nM PA preincubate with each other for 15 minutes before the addition of RNA substrate. B. The inhibitory effect of various concentrations of BXA on the endonuclease activity of H1N1 PAc. The results were quantified from three replicates. Various concentrations of BXA and 300nM PAc preincubate with each other for 15 minutes before the addition of RNA substrate. C. The structure of H17N10 PA with the interaction of BXA. PA was shown in pink and BXA molecule was shown in green. Predicted interaction sites are highlighted in red. D. Amino acid alignment of H17N10 and several important human influenza A virus PA. The key residues possibly important for the interaction between PA and BXA as indicated in fig3C was highlighted. E. Endonuclease activity of H1N1 wild-type and mutated PA. Residues that may be critical for the interaction of H1N1 PA and BXA were mutated into Ala. Equal amounts of wild-type and mutant proteins were applied to RNA degradation reactions to compare their endonuclease activities. F. Inhibitory effect of BXA on H1N1 wild-type PA and F707A mutant. 1μM BXA and 300nM wild-type PA or F707A mutant preincubate with each other for 15 minutes before addition to the RNA degradation reactions. G. Quantitative result of fig3F from three replicates. Image J software was used to analyze the product bands to quantify the enzyme activity. ****: p < 0.0001.

### Screening of anti-influenza compounds by targeting the PAc domain

Confirmation of novel antiviral drug targets in the PAc domain of the influenza virus could provide new insights related to drug screening. To narrow down the potential compounds, virtual screening was performed to identify those chemicals with higher binding affinity for the PAc domain (Fig. 4A). Among 300,000 compounds screened, 1,000 showed higher docking scores. Forty-six compounds were selected to further detect the antiviral activity in MDCK cells. Methotrexate (MTX) and A-1155463 exhibited a higher anti-influenza virus effect at a concentration of 10 μM (Fig. 4B). Because A-1155463 was found to be clearly cytotoxic, MTX was investigated further (Fig. S6). With treatment using various concentrations of MTX, the half-maximal effective concentration (EC_50_) value was calculated to be 12 nM, and the cytotoxic concentration 50% (CC_50_) value was more than 25 μM (Fig. 4C). The high antiviral efficiency and low cytotoxicity indicated that MTX is a novel drug against the influenza virus. Compared with influenza virus-infected cells, cells treated with MTX showed significantly decreased levels of viral PA and nucleoprotein (Fig. 4D). Immunofluorescence staining targeting the PA protein also demonstrated that MTX can inhibit the replication of influenza virus (Fig. 4E). Molecular docking between MTX and PA indicated the binding sites in the protein (Fig. 4F and Fig. S7). With the addition of varying amounts of MTX to the PA endonuclease reaction system, the enzyme activity of PA was suppressed with increased concentrations of MTX (Fig. 4G). Similarly, as analogs of MTX, aminopterin and pralatrexate also inhibit the replication of the influenza virus at concentrations of the nM scale (Fig. S8). Folic acid supplement had no obvious effect on the antiviral activity of MTX (Fig. S9). These results proved that MTX inhibits influenza virus replication by targeting the PAc domain, which suppresses PA endonuclease activity.

**Fig. 4.**
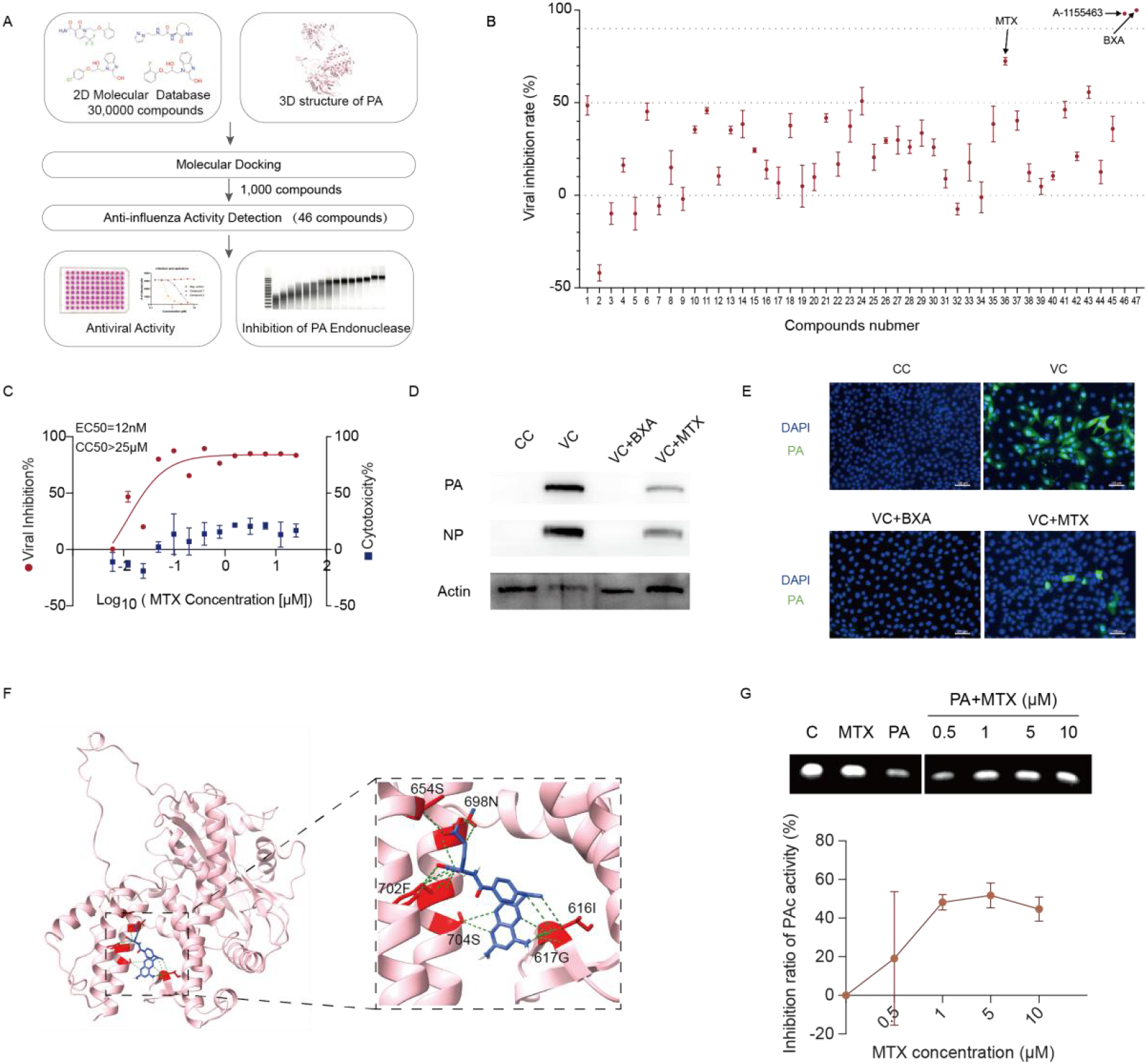
MTX was discovered in high-throughput screening targeting the influenza PAc domain. A. Screening workflow to identify antiviral compounds targeting PAc domain. A virtual screening of 300,000 compounds against the PA protein yielded approximately 1,000 high-scoring candidates based on docking scores, with 46 compounds applied in subsequent functional assays to identify the potential antiviral compounds. B. Anti-influenza effect screening of 46 compounds in MDCK cells at a concentration of 10 µM. BXA was used as the positive-drug control. C. EC_50_ and CC_50_ measurements of MTX in MDCK cells. The EC_50_ of the compounds was determined by measuring the viral titer in the supernatant of MDCK cells treated with virus and compounds at various concentrations for 24 hours. CC_50_ of compounds was assessed by using CCK8 kit according to the instruction manual. D. MDCK cells were infected with H1N1, with or without the treatment of MTX. Western blot was used to detect the viral PA and NP levels in the cell lysates. Actin was used as a reference gene, and BXA was used as the positive drug control. E. Immunofluorescence staining was used to detect influenza PA in MDCK cells after infection with H1N1 virus for 24h with or without the addition of MTX. BXA was used as the positive drug control. Cell nucleus was stained with DAPI. Scale bar: 100 μm. F. Structure of influenza PA with the interaction of MTX. MTX molecule was in blue color and PA protein was in pink color. The potential key residues that vital for the interaction were highlighted in red. G. Inhibitory effect of various concentrations of MTX on the endonuclease activity of PA. Various concentrations of MTX and 300nM PA preincubate with each other for 15 minutes before addition to the RNA degradation reaction system. The results were quantified from three replicates. Image J software was used to analyze the product bands to quantify the enzyme activity.

### MTX treatment suppresses influenza virus propagation in vivo

To evaluate the therapeutic effect of MTX, we administered the drug at different time points post-infection, as indicated. Lung tissue from the experimental group was collected at 4 days post-infection for further analysis (Fig. 5A). By monitoring body weight, normal mice and the MTX-only control group exhibited continuous weight gain, suggesting that the administered dose of MTX did not cause significant toxicity. Compared with the virus-infected group, the MTX treatment group showed a reduced degree of weight loss (Fig. 5B). Moreover, mice who received treatment with MTX had a longer survival duration after infection with the influenza virus (Fig. 5C). These findings demonstrate that MTX confers a therapeutic benefit in severe influenza infection. To further investigate the antiviral effects of MTX, we quantified the viral load in mouse lung tissue and found that the virus content in the MTX treatment group was significantly lower than that in the virus-infected group (Fig. 5D). In addition, immunohistochemistry of lung tissue showed lower influenza PA protein levels in the MTX treatment group, which signified that MTX can effectively inhibit viral replication in lung tissue (Fig. 5E). MTX treatment significantly alleviated lung injury on pathological examination (Fig. 5F and Fig. S10). With MTX treatment, inflammatory infiltration, alveolar wall thickening, alveolar expansion, eosinophilic material, and inflammatory exudation were notably decreased. These results demonstrated that MTX can effectively inhibit influenza virus reproduction and alleviate symptoms of infection in vivo in a mouse model.

**Fig. 5.**
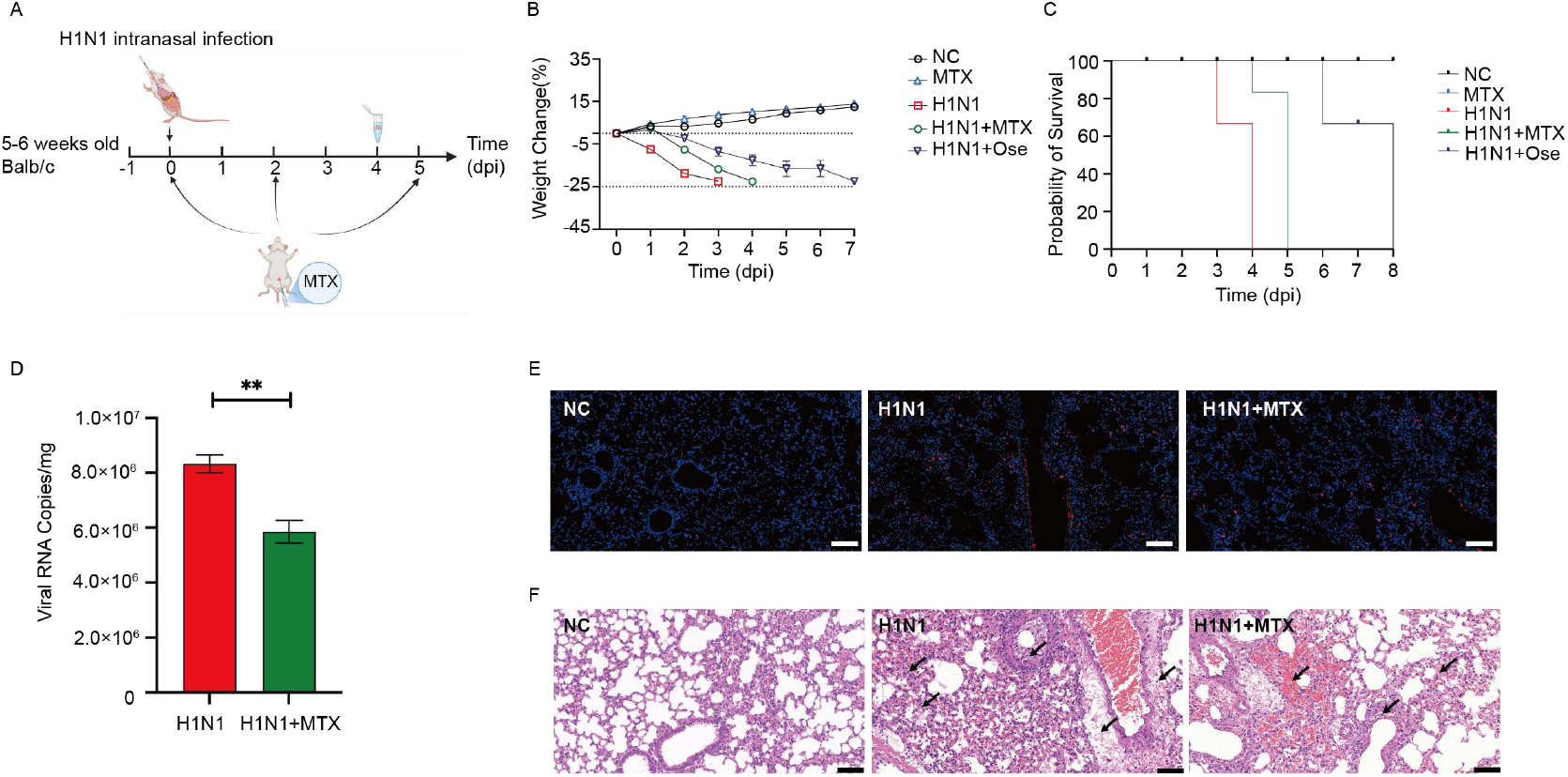
Therapeutic effects of MTX on severe influenza-infected mice. A. Experimental timeline of MTX treatment in influenza-infected mice. MTX was administered by intraperitoneal injection at a dose of 13.5 mg/kg/day on the day of infection, 2 dpi, and 5 dpi. Oseltamivir was administered once daily via oral gavage at a dose of 60 mg/kg/day. Lung tissue was collected at 4 dpi. B. Body weight changes of mice in normal, MTX-only, virus-infected, MTX-treated and oseltamivir-treated groups. Values are means, with their standard errors represented by vertical bars. C. Survival curve of mice in normal, MTX-only, virus-infected, MTX-treated and oseltamivir-treated groups. D. Viral load quantification in lung tissues of mice in virus-infected and MTX-treated groups. RNA was extracted from mouse lungs that collected at 4 dpi and RT-qPCR was used to quantify the viral load. ***: p < 0.001vs Virus control group (H1N1). E. Immunofluorescence staining of mice lung tissues in the normal, virus-infected and MTX-treated groups. Influenza PA protein was indicated in red that stained by influenza PA antibody and fluorescent secondary antibody. Cell nucleus was in blue by DAPI dye. Scale bar: 50 μm. F. Pathological examination of mice lung tissues in the normal, virus-infected and MTX-treated groups by HE staining. The black arrows indicate the presence of hyperaemia and infiltration of inflammatory cells in the tissues. Scale bar: 50 μm.

## Discussion

Our work fundamentally challenges the conventional functional domain paradigm of PA protein and redefines the framework for future mechanistic investigations. As the long-standing consensus, the endonuclease active site of influenza virus PA protein resides within its N-terminal domain (*19, 20*). This study demonstrates through systematic experimentation that PAc retains high endonuclease activity, whereas PAn exhibits minimal catalytic capacity under 10-minute reaction conditions. Of note, previously reported PAn with enzyme activity is located within residues 1-209, while its catalytic process generally requires a reaction duration of over 60 minutes. Therefore, PAc was firstly identified to be the bona fide active center of PA with high endonuclease activity, which enabled us to further explore the viral RNA replication mechanism. Y393, located in the C-terminal of H1N1 PA, is reported to be indispensable for viral RNA amplification, which is consistent with our research (*22*). In our study, crystallographic structural analysis identifies critical RNA-binding residues (YDS motif at positions 393-395) within the PAc domain, with mutagenesis experiments confirming the indispensable role of these residues in catalytic function. These structural insights establish a revised molecular basis for understanding PA-mediated endonuclease activity and its regulatory mechanism.

Baloxavir acid (BXA), a first-line anti-influenza drug approved in recent years, requires further clarification regarding its precise molecular target (*23, 24*). Through functional assays and molecular docking, this study definitively demonstrates that PAc, not PAn serves as the direct binding site for BXA. This critical finding establishes a structural foundation for optimizing BXA and its derivatives to enhance potency and selectivity. Furthermore, we identified that the F707A mutation within the PAc domain abolishes BXA-mediated inhibition, indicating that residue F707 likely represents a high-risk hotspot for clinical resistance mutations. This discovery underscores the necessity of monitoring F707 alterations in global influenza surveillance programs and highlights its predictive value for guiding next-generation antiviral development (*25*).

Our results provide a novel strategy for anti-influenza drug development by targeting PAc. Through virtual screening against PAc combined with antiviral assays, we identified methotrexate and aminopterin as potent antiviral compounds with EC_50_ values at the nanomolar level. These clinically approved agents, currently used for treating malignancies and autoimmune disorders, demonstrate the potential of drug repurposing strategies to accelerate clinical translation of influenza therapeutics (*26*-*29*). Notably, methotrexate exhibited significant antiviral efficacy in both cellular and murine infection models, effectively suppressing viral replication and alleviating disease symptoms. Our findings validate PAc as a viable antiviral target and establish a foundation for optimizing existing compounds or developing novel PAc inhibitors through structure-based drug design approaches.

The potential clinical significance was indicated through dual translational perspectives. On one hand, compounds such as methotrexate may exhibit synergistic antiviral effects when combined with existing influenza therapeutics, offering novel combinatorial regimens for managing severe influenza cases. On the other hand, the development of PAc-targeted antivirals could address unmet clinical needs caused by BXA-resistant strains, particularly through rational drug design strategies targeting critical resistance-associated mutations such as F707. Furthermore, the integration of cross-disciplinary methodologies—including structural biology (crystallographic studies), biochemistry (mutational profiling), computational biology (molecular docking), and pharmacology (in vitro and in vivo validation)—not only validates PAc as a druggable target but also provides a methodological reference for analogous antiviral research.

This study has several limitations that warrant further investigation. First, the precise role of PAc in the influenza viral replication cycle requires deeper mechanistic elucidation, particularly regarding its cooperative mechanisms with other polymerase subunits (e.g., PB1 and PB2). Second, the functional interplay between the PAn and PAc domains remains unresolved; whether they operate through synergistic or complementary interactions demands systematic exploration. Third, while methotrexate demonstrates promising antiviral activity, its immunosuppressive properties raise safety concerns for prolonged influenza treatment, necessitating structural optimization to mitigate off-target effects. To advance preclinical development, future work should validate therapeutic efficacy in physiologically relevant models (e.g., ferret infection systems) and assess the emergence risk of drug-resistant mutants.

In summary, our work not only advances the mechanistic understanding of influenza PA protein functionality but also pioneers innovative avenues for antiviral development through target repositioning and drug repurposing strategies. Its scientific significance lies in correcting long-standing misconceptions while providing translatable therapeutic prospects, particularly in addressing influenza drug resistance and emerging strains. Future investigations should prioritize structure-guided optimization of PAc inhibitors, discovery of novel scaffolds targeting PAc, and systematic evaluation of combination therapeutic regimens to maximize clinical efficacy and resilience against viral evolution.

## Acknowledgement

We thank professor Lei Shi in Peking Union Medical College for the scientific suggestions. And we also thank Dr. Zhiguo Mang in Guizhou medicine Law natural medicine Technology Co., LTD for assisting in structure analysis of compounds.

## Author contributions

Conceptualization: S.X., Y.G.T., W.W.; Funding acquisition: Y.G.T., S.X.; Investigation: S.X., X.Y.W., L.H.Z., Y.L.T., Y.Q.W., S.Y.D.; Methodology: S.X., X.Y.W., L.H.Z., Y.L.T.; Supervision: Y.G.T., S.X., W.W.; Visualization: S.X., X.Y.W., L.H.Z.; Writing – original draft: S.X., X.Y.W., L.H.Z.; Writing – review & editing: Y.G.T., W.W., S.X., X.Y.W., L.H.Z., Y.L.T. Competing interests: All authors declare that they have no competing interests.

## Data and materials availability

All data are available in the paper or in the supplementary materials.

## Supplementary Materials

Materials and Methods

Figs. S1 to S10

