## supplementary for "Identification of the bona fide active center of influenza A virus polymerase acidic protein as the antiviral target"

### Materials and Methods

#### Plasmids construction

The coding sequence of H1N1 PA was optimized to enhance the expression in *Escherichia coli* (*E.coli*). The optimized PA, PA N-terminal (PAn)and PA C-terminal(PAc) sequences were cloned into the *Nde*I and *Xho*I restriction sites of pET28a vector. PA and PAc mutants were obtained by site-directed mutagenesis according to the manual of Mut Express II Fast Mutagenesis Kit V2 (C214,Vazyme,) .All constructed plasmids were confirmed by first-generation sequencing.

#### Proteins expression and purification

The plasmid containing the target gene was transformed into *E.coli* BL21 (DE3) cells. BL21 cells were cultured in Luria-Bertani (LB) medium containing 50 μg/mL kanamycin at 37℃ and 200 rpm until the $OD_{600}$ was 0.8. IPTG was then added to a final concentration of 0.4 mmol/L to induce the protein expression at 16℃ and 150 rpm for 24 h. After collected by centrifugation at 6000 rpm for 10minutes, bacteria were resuspended in lysis buffer (20 mmol/L Tris-HCl, 500 mmol/L NaCl, 0.5% TritonX-100, 20 mg/L RNaseA, 20 mg/L DNaseI, 10% (v/v) glycerol, 1 mmol/L DTT, 1×protease inhibitors, pH=8.0), and disrupted in a cell crusher. Cell debris was removed by centrifugation at 18000rpm for 45 minutes at 4℃. The Ni-NTA Agarose was added to the supernatant to bind the target protein at 4℃ for 4h. The Ni-NTA column was washed twice with wash buffer (20 mmol/L Tris-HCl, 500 mmol/L NaCl, 0.4 mmol/L DTT, and 30 mmol/L imidazole, pH=8.0), followed by wash buffer with higher imidazole concentration (20 mmol/L Tris-HCl, 500 mmol/L NaCl, 0.4 mmol/L DTT, 60 mmol/L imidazole, pH=8.0). Elution buffer (20 mmol/L Tris-HCl, 500 mmol/L NaCl, 500 mmol/L imidazole,1 mmol/L DTT, 10% glycerol, pH=8.0) was added to collect target proteins. After concentrated by 30 KD centrifugal filters, proteins were dialyzed in dialysis buffer (20 mmol/L Tris-HCl, 100 mmol/L NaCl, 10% glycerol, and 1 mmol/L DTT, pH=8.0). Purified proteins were detected by Coomassie brilliant blue staining in SDS-PAGE gel. Proteins were flash frozen using liquid nitrogen and stored at −80℃.

PA endonuclease activity detection

ph-RNA (5'-AGUAGAAACAAGGGUAUUUUUCUUGGGACGCCAUGAUUUUGAUGUCACUCAGUG AGUGAUUAUCUACCCUGCUUUUGCU-3') was the substrate of PA as was reported (19). 750nM ph-RNA was co-incubated with 300nM PA protein and 1 mM $MnCl_2$ in reaction buffer solution (20 mM Tris-HCl, 100 mM NaCl, pH=8.0) for 10 minutes at 37℃, and then EGTA with a final concentration of 20 mM was added to stop the reaction. To test the effect of compounds on endonuclease activity, compounds and protein preincubate with each other for 15 minitutes before the addition of RNA substrate. The reaction products were detected by electrophoresis on an 8% urea-polyacrylamide gel and staining with YeaRed Nucleic Acid Gel Stain.

For fluorescence resonance energy transfer (FRET) assay, 750 nM of fluorescently labeled RNA (6-FAM-5'-CUCCUCAUUUUUCCCUAGUU-3'-BHQ1) was incubated with the protein in the same buffer as previously described for the endonuclease activity assay. The fluorescence intensity was subsequently measured using a BioTek Synergy H1 microplate reader at room temperature, with an excitation wavelength of 465 nm and an emission wavelength of 520 nm.

Virtual screening

Protein Preparation Wizard module was employed to add hydrogen atoms to PA structure in H17N10 RdRp complex ( PDB:6t0n ), followed by energy optimization using the OPLS2005 force field with a root mean square deviation (RMSD) of 0.30 Å. The docking pocket was defined as a region encompassing the three key residues (Ile623, Lys638, and Phe705). The dimensions of the grid box were set to 20 Å × 20 × 20 Å. A total of 300,000 compounds in 2D format were processed using Schrodinger's LigPrep Module for hydrogenation and energy optimization; subsequently, their corresponding 3D structures were generated for virtual screening. Virtual Screening Workflow module was used to perform the virtual screening by importing prepared compounds and employing Glide for molecular docking. Specifically, PA and compound molecules underwent geometric matching and energy matching during docking procedures. The library compounds were docked against a newly identified active pocket on PA protein utilizing high throughput virtual screening (HTVS) mode. From this initial round, we selected the top 10% of small molecule candidates for further evaluation through standard precision (SP) mode screening. Subsequently, we refined our selection by choosing another top 10% based on scoring values in high precision (XP) mode to ultimately rank small molecular compounds. The results from docking studies were visualized using ChimeraX software, allowing us to analyze key amino acid sites along with their interaction forces.

Cell culture and virus infection

Madin-Darby canine kidney (MDCK) cells were grown in Dulbecco's Modified Eagle's Medium (DMEM/HIGH GLUCOSE, HyClone), supplemented with 10% fetal bovine serum (FBS, PAN-Biotech) and 1% penicillin/streptomycin (HyClone) at 37℃ with 5% $CO_2$. H1N1 virus (A/PR/8/34, ATCC, VR-1469) and H3N2 virus (A/Aichi/2/68, ATCC, VR-1680) was utilized in this study. The viral stock was diluted in phosphate-buffered saline (PBS) containing 0.2% bovine serum albumin (BSA) with a small volume, and incubated with MDCK cells for 2 hours at 37℃ in a humidified atmosphere of 5% $CO_2$. Following the incubation period, DMEM medium supplemented with 2 μg/mL TPCK-trypsin, 1% penicillin/streptomycin, and an additional 0.2% BSA was added to the culture and incubated for a further duration of 48 hours. After incubation, the supernatant was collected and centrifuged at 3000 rpm for 10 minutes at a temperature of 4℃to eliminate cell debris. The resulting supernatant was then collected, aliquoted into liquid nitrogen for flash freezing, and subsequently stored at -80℃.

Western blot

MDCK cells were infected with H1N1 virus (200 $TCID_{50}$/mL) in the presence of BXA at a concentration of 100nM or methotrexate at concentrations of 1 μM. 24 hours after infection, cell samples were lysed in RIPA buffer to detect virus PA and NP protein levels by Western blot. Influenza A virus PA protein antibody (GTX636828, GeneTex), influenza A virus nucleoprotein antibody (GTX125989, GeneTex) and β-Actin antibody (ABL1010， Abbkine)were used as primary antibodies. HRP,Goat Anti-Rabbit IgG (A21020, Abbkine) or HRP, Goat Anti-Mouse IgG (A21010, Abbkine) were used as secondary antibodies.

Immunofluorescence

MDCK cells were seeded on glass slides in 24-well plates and infected by H1N1 virus with the treatment of different compounds. 24 hours after infection, cells were fixed with 4% paraformaldehyde for 10 minutes 24h after infection. After fixation, cells were permeabilized with 0.2% Triton X-100 in PBS for 10 minutes at room temperature. Cells were subsequently blocked in PBS containing 3% BSA for 30 minutes at room temperature and incubated with PA antibodies (GTX636828, GeneTex) for 2h at 37℃. After washing three times with PBS, Dylight 488, Goat Anti-Rabbit IgG (A23220, Abbkine) was as secondary antibody and incubated for 1 h at 37℃. After washing three times with PBS and one time with $ddH_2O$, the samples were stained with DAPI by Antifade Mounting Medium (S2110,Solarbio) and subsequently viewed by fluorescence confocal microscopy.

Assessment of $EC_{50}$ and $CC_{50}$ of compounds

MDCK cells was seeded in 96-well plates before incubating with influenza virus (1000 $TCID_{50}$/mL) and various concentrations of compounds. 1 hour after incubation, cells were washed by PBS and then cultured in DMEM medium supplemented with 2 μg/mL TPCK-trypsin, 1% penicillin/streptomycin, 0.2% BSA and compounds for 24h. Cell supernatant was collected and total viral RNA was extracted using the Super FastPure Cell RNA Isolation Kit (RC102-01, Vazyme). vRNA was reverse transcribed into cDNA using HiScript II Q RT SuperMix for qPCR (+gDNA wiper) (R223-01, Vazyme). qPCR was performed using QuantStudio 1 by Taq Pro Universal SYBR qPCR Master Mix (Q712-02, Vazyme) to assess the virus titration. The primer sequences were as follows: H1N1-F (5’-GACCAATCCTGTCACCTCTGAC-3’), H1N1-R (5’-GGGCATTTTGGACAAAGCGTCTACG-3’).

The viral inhibition rate (VIR) was calculated as followed: VIR=(viral RNA copies in virus alone group - viral RNA copies in compounds group)/viral RNA copies in virus alone group. GraphPad-Prism was used to calculate $EC_{50}$ of compounds. Cytotoxic effect of compounds was assessed by using CCK8 kit (BS350C, Biosharp) according to the the instruction manual. CC50 value was calculated by GraphPad-Prism.

Antiviral effect in vivo

Five to six weeks old SPF BALB/c mice weighing 15-18 g were purchased from Beijing Vital River Laboratory Animal Technology Co., Ltd. The mice were housed in individually ventilated cages (IVCs), with access to sterile water and sterile mouse feed provided ad libitum. $LD_{50}$ of H1N1 was determined to be 2×10^4.5 $TCID_{50}$/mL and 100 $LD_{50}$ was used in vivo. Oseltamivir was administered once daily via oral gavage using a plastic feeding tube (22 ga, black, × 25 mm) at a dose of 60 mg/kg/day, starting the day before virus infection and continuing until 5 dpi. MTX was administered by intraperitoneal injection at a dose of 13.5 mg/kg/day on the day of infection, 2 dpi, and 5 dpi. Lung tissues were collected on 4 dpi for subsequent experiments.Typically, 6 mice were used for administration of each dose during viral challenge to detect the weight change and survival rates. RNA was extracted from mouse lungs using TRIzol reagent (Invitrogen, Carlsbad, CA, USA). cDNA was synthesized using the PrimeScript™ RT Reagent Kit with gDNA Eraser (Perfect Real Time) (TAKARA, Japan). qPCR method was as the mentioned above. Lung tissues were collected and fixed in 10% formalin for 72 hours, followed by dehydration with a graded ethanol series and paraffin embedding. Tissue sections (5 μm thick) were stained with hematoxylin and eosin (H&E) to evaluate histopathological changes. Immunohistochemistry of lung tissue paraffin sections was performed as routing procedure by using influenza PA antibody (GTX636828, GeneTex).

Ethics statement

All animal procedures were conducted under protocols approved by the Animal Care and Use Committee of Ocean University of China (OUC-SMP-2024-07-02).

Statistical analysis

All data are representative of at least three independent experiments and expressed as the mean ±SD. The data was analyzed with the GraphPad Prism 9 software to evaluate the statistical significance by two-tailed unpaired Student's t-test. A statistically significant difference was accessed when $P < 0.05$. $P < 0.05$ was marked *. $P < 0.01$ was marked **, $P < 0.001$ was marked *** and $P < 0.0001$ was marked ****.

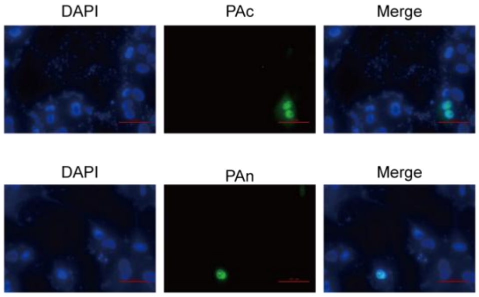


Fig.S1. PAc has a specific subcellular localization.Immunofluorescence staining was used to detect PAc and PAn in MDCK cells 24h after infection with H1N1 virus. Scale bar: 100 μm.

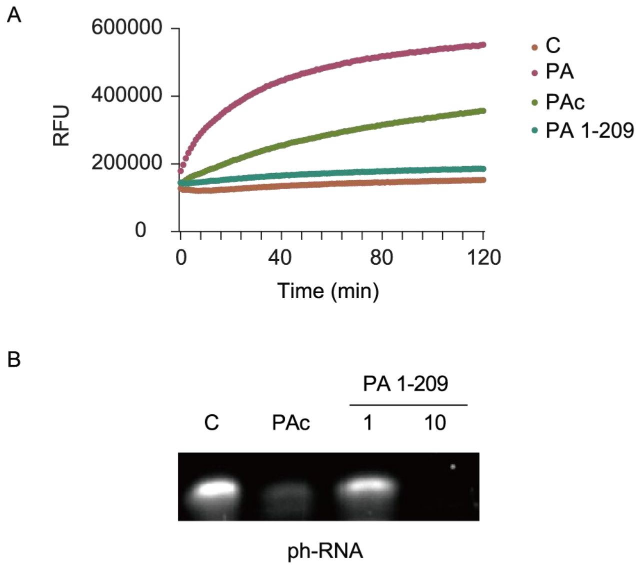


Fig.S2. The endonuclease activity of PAc was higher than that of the PA 1-209 truncate. A. The FRET-based assay was used to detect the endonuclease activities of PA, PAc and PA 1-209 with FAM/BHQ1-labeled RNA as the substrate. B. Endonuclease activity of PAc and PA 1-209 (at concentrations equivalent to and 10-fold higher than PAc) with RNA as substrate. The reaction time was 60 minutes.

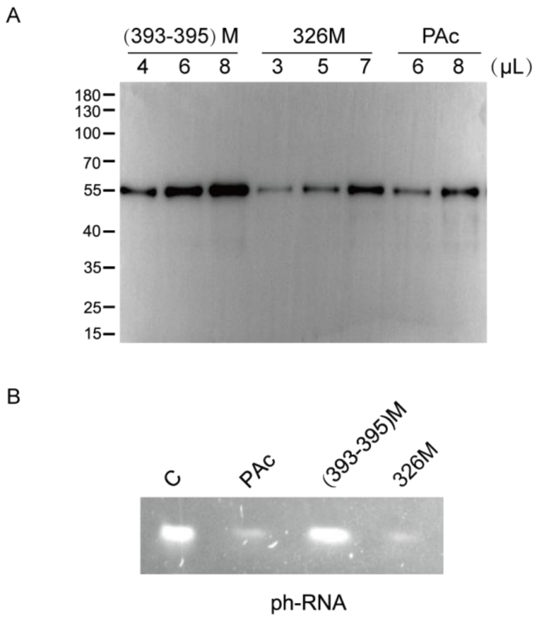


Fig.S3. Semi-quantitative and activity comparison between wide-type and mutated PAc. A. Purified (393-395) mutant, 326 mutant and PAc were semi-quantified by western blot using anti-His tag antibody. B. Endonuclease activity of PAc, (393-395) mutant and 326 mutant with RNA as substrate.

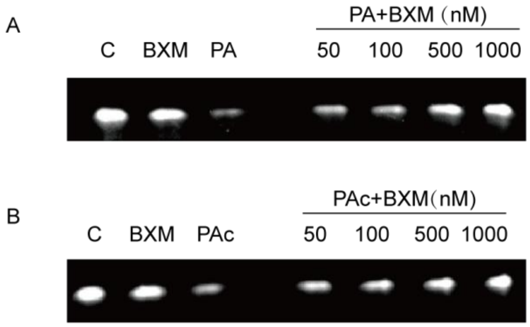


Fig.S4. Effect of BXM on the enzyme activity of PA and PAc. A. The inhibitory effect of various concentrations of BXM on endonuclease activity of H1N1 PA. B. The inhibitory effect of various concentrations of BXM on the endonuclease activity of H1N1 PAc.

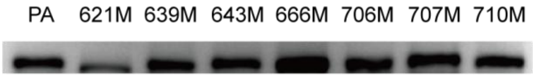


Fig.S5. H1N1 wild-type and mutated PA were semi-quantified by western blot using Influenza A virus PA protein antibody.

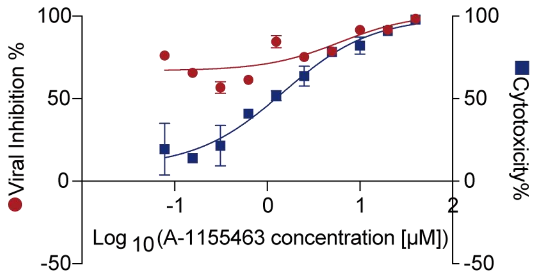


Fig.S6. The anti-influenza effect and cytotoxicity of A-1155463 in MDCK cell. The antiviral effect of the drug was determined by measuring the viral titer in the supernatant of MDCK cells 24h after infection with H1N1 virus. Cytotoxicity of compounds was assessed using CCK8 kit according to the instructions.

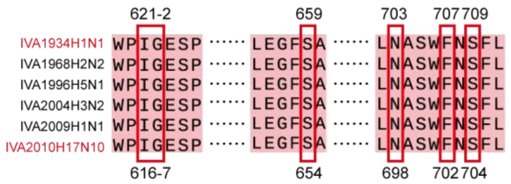


Fig.S7. Amino acids alignment of H17N10 and some human influenza A PA. The key residues that possibly important for the interaction of PA and MTX were indicated.

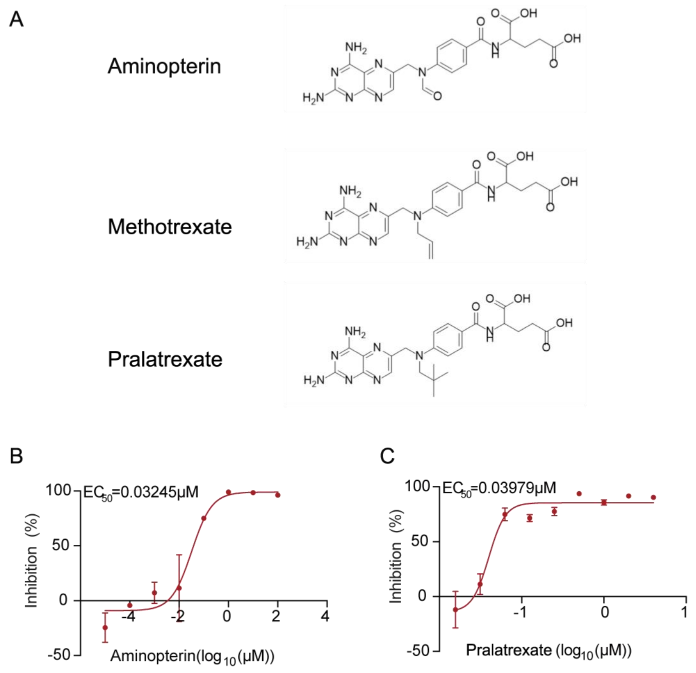


Fig.S8. The methotrexate analogues aminopterin and pralatrexate effectively inhibit influenza virus replication. A. Chemical structural formulae of aminopterin, methotrexate and pralatrexate. B. $EC_{50}$ measurement of aminopterin in MDCK cells. C. $EC_{50}$ measurement of pralatrexate in MDCK cells.

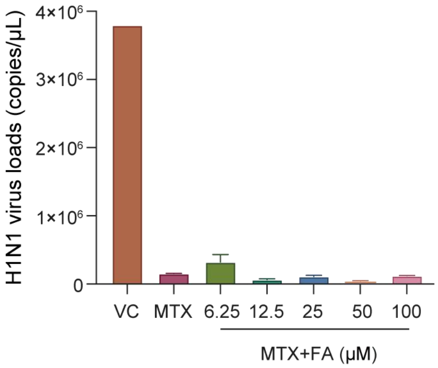


Fig.S9. Antiviral effects of methotrexate are not related to folic acid. MDCK cells were infected with H1N1 virus and treated with 1 μM MTX alone or in combination with increasing concentrations of FA (6.25, 12.5, 25, 50, and 100 μM). After 24 hours, viral RNA levels were quantified by qPCR. The MTX-only group served as a reference to evaluate whether FA supplementation alters the antiviral effect of MTX. Data represent mean ± SD from three independent experiments.

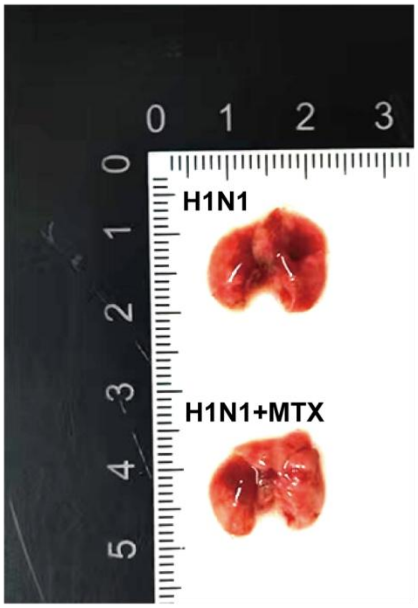


Fig.S10. Representative excised lungs from H1N1-infected mice.Top:Severe H1N1-induced lung damage (extensive hemorrhage).Bottom:Attenuated damage in H1N1-infected mice treated with MTX.
